# Detection of protein-protein interactions by bio-orthogonal fluorogenic proximity probes

**DOI:** 10.1101/2024.08.20.608485

**Authors:** Andreas Torell, N. Alfred Larsson, Storm Phillipson, Luke R. Odell, Daniel Fürth

## Abstract

Detecting protein-protein interactions within cells is challenging. Transgenic approaches risk altering protein function via fluorescent tagging, while *in situ* methods lack *in vivo* compatibility. Here, we introduce fluorogenic probes with dual-tetrazine pegylated branched arms linked to xanthene dye. Activation requires both tetrazine arms to interact simultaneously with target proteins, enabling dual-substrate recognition. We applied our method to detect protein-protein interactions in both fixed and living cells, utilizing antibody conjugation for fixed cells and genetic code expansion for real-time detection in living cells. Our strategy ensures versatile applicability and seamless transition between fixed and living systems.

## Introduction

Numerous techniques exist for detecting protein-protein interactions and determining their cellular locations. These approaches can be broadly categorized into two main groups: transgenic fusion-protein techniques (1, 2), and methods employing a combination of affinity reagents with either fluorescence resonance energy transfer or enzymatic detection and amplification techniques using oligonucleotides for readout (3–5). Transgenic techniques offer the distinct advantage of enabling live imaging, allowing for real-time monitoring of interactions within living cells (6). In contrast, the latter category of methods, utilizing affinity reagents and enzymatic amplification techniques, excels in detecting endogenous proteins where transgenic approaches are not feasible, such as in clinical samples (3).

What has been lacking is a fluorescent technique that operates independently of transgenic expression of fusion proteins, thus sidestepping potential disruptions in protein function due to tagging, while also addressing sensitivity and reliability challenges typically associated with multi-step enzymatic detection and amplification procedures. A unified detection method capable of functioning effectively in both fixed and living cells would bridge these two prominent methodological categories.

In tandem with these challenges, proximity labeling techniques, despite their achievements (7–9), face a distinct issue. A persistent challenge lies in defining a precise spatial distance that signifies interaction between proteins. This limitation stems from the reliance on proximity-driven events without a well-defined threshold for spatial constraints. The ambiguity in interpreting proximity labeling data mirrors the limitations of traditional pulldown assays, posing an impediment to precisely delineating the spatial proximity necessary for accurate molecular interactions and contributing to the occurrence of false positives.

Here we address these dual challenges, we provide a comprehensive solution that not only operates independently of transgenic expression but also offers precise control of spatial distances, akin to a ‘molecular ruler’—a mechanism that enables the accurate measurement of distances between biomolecules. The integration of non-enzymatic and bio-orthogonal ligation chemistries with fluorogenic properties presents a promising solution to fulfill this critical need (10–12). To this end, we developed and validated a fluorogenic dual-substrate recognition probe. We investigate its specificity in detecting protein-protein interactions, highlighting its capability to distinguish true interactions from mere proximity detection. Furthermore, we employ our probes for the detection of endogenous targets using conjugated antibodies and real-time monitoring of dynamic protein-protein interactions in live cells using genetic code expansion.

## Fluorogenic Proximity Probes (FluoroProx)

Our method involves the synthesis of fluorogenic probes, linking a xanthene-derivative dye to a branched dual-substrate recognition arm through poly(ethylene glycol) (PEG) linkers. The fluorescence of these probes only activates upon the simultaneous recognition of both arms by their respective targets. Branched PEG linker arms have long enabled researchers to synthesize compounds with multiple terminal functional groups generating superior properties in cellular uptake and biodistribution (13, 14). These arms are structured to position two reactive groups at a precise distance from each other (**Fig. S1a**), facilitating proximity-driven transformation of the second reactive group when brought into close proximity by the reaction of the first. We utilized branched PEGylation of a fluorophore to bring into close proximity two tetrazines (**Figure 1a**). Tetrazines in these fluorogenic probes play a dual role. Firstly, they serve as diene groups, engaging in an inverse electron-demand Diels–Alder reaction with their target substrate containing a dienophile (15). Secondly, tetrazines also function as fluorescence quenchers for the fluorophore (**Fig. 1b**). The fluorophore can thus exist in three different states-a fully quenched state, a semi-quenched state and an unquenched state (**Fig. 1a**), corresponding to no antigen detection, monomer detection, and dimer detection. The probes are easily synthesized with common molecular biology equipment through strain-promoted azide-alkyne cycloaddition conjugation of dibenzocyclooctyne-containing fluorophores, yielding a high product yield and eliminating the need for subsequent purification (**Fig. S1a,c**).

**Figure 1.**
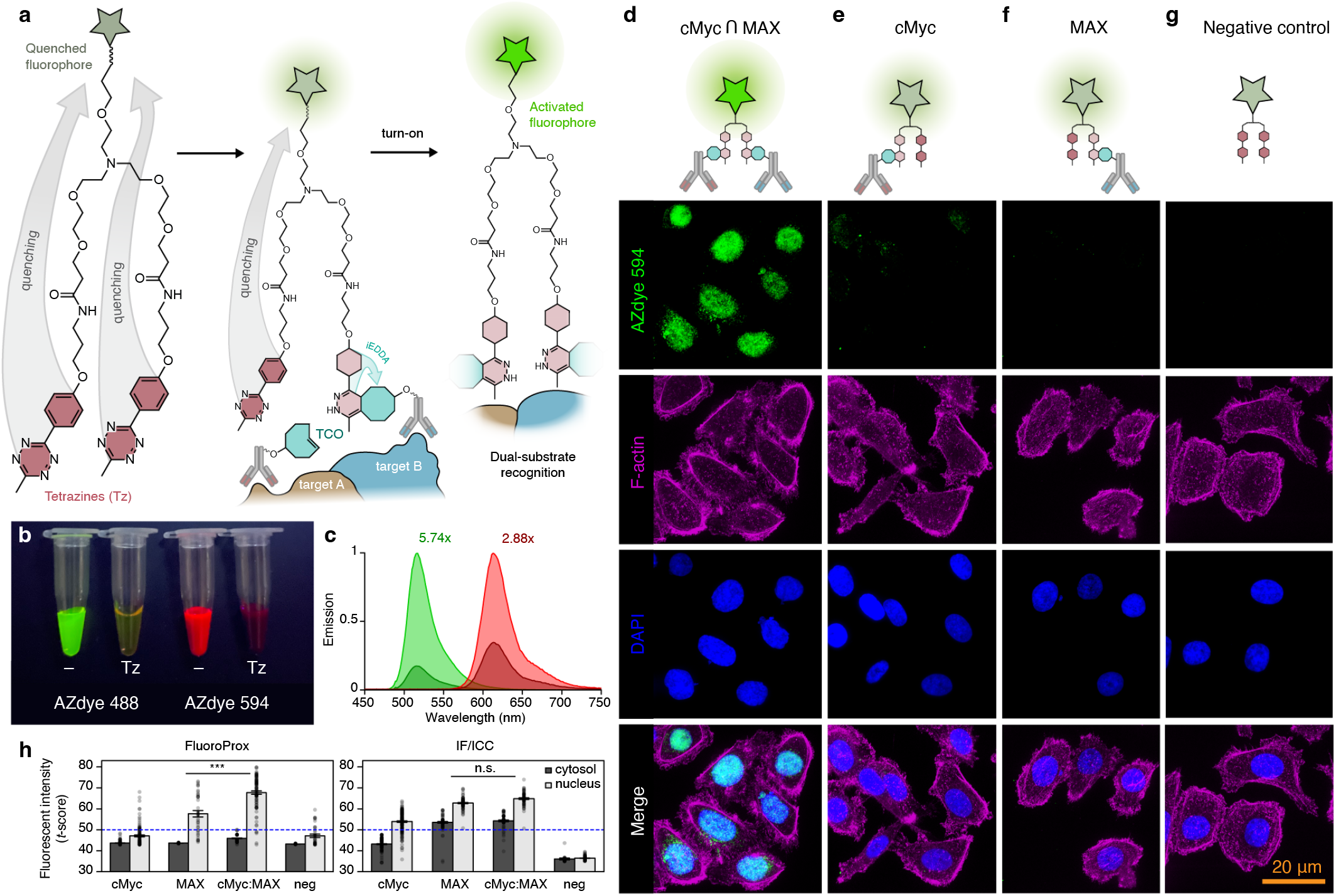
Fluorogenic Proximity (FluoroProx) Probes Enable Dual-Substrate Recognition. **a)** Schematic of fluorogenic proximity probes. A branched PEG linker with three arms connects a xanthene-derivative dye to two tetrazines. The energy transfer and quenching of the dye are mitigated only upon reaction of the tetrazine with trans-cyclooctene conjugated antibodies, resulting in rapid fluorogenic activation. **b)** Quenching of AZDye488 and AZDye594 upon conjugation to PEG1-bis(PEG2-tetrazine), Tz, compared to unquenched, -. **c)** Emission spectra of AZdye488 and AZdye594 before and after bis-tetrazine quenching. **d)** Detection of the cMyc/MAX transcription factor oncogene complex in PC3 cells. Primary cMyc and Max antibodies are detected by secondary antibodies conjugated to trans-cycloctene (TCO), which serves as the substrate for the branched tetrazine arm’s inverse electron-demand reaction. Dual-substrate recognition results in complete unquenching of the fluorescent signal. **e)** Single primary antibody presence, either cMyc or **f)** Max, is insufficient to reverse the quenching effect caused by unreacted free tetrazine arms. **g)** Omission of secondary TCO-antibodies. Green: AZDye594 signal, magenta: Phalloidin-FITC, blue: DAPI. Scale bar: 20 μm. In the schematics of d-g, the indirect antibody staining is omitted for simplicity. **h)** Quantitative comparison of fluorescent signal with FluoroProx probes compared to traditional immunofluorescence for cMyc/MAX in nucleus versus cytosol. Groups indicate presence of primary antibody. Neg., are negative controls, where all primary antibodies were ommitted, only secondary used (*** *P* < 0.001, n.s. *P* > 0.05). Error bars +/-1 S.E.M. *n* = 689 cells

Prior studies have established that quenching efficiency of tetrazine dyes is significantly affected by the distance between the tetrazine and fluorophore chromophores (16). Consequently, our initial investigation aimed to assess whether the PEG-linker of our probes introduced an excessive distance between these chromophores, potentially compromising fluorescent turn-on for practical applications. We evaluated quenching of both a green-fluorescent dye (AZDye488, λ_abs_ 490, λ_em_ 517 nm) as well as a red-fluorescent dye (AZDye594, λ_abs_ 590 nm, λ_em_ 613 nm). Despite the increased distance introduced by the PEG-arm, both dyes consistently demonstrate potent quenching capabilities (**Fig. 1b**), yielding fluorescent quench emission ratios of 5.74 and 2.88, respectively (**Fig. 1c**). This highlights the system’s capability to effectively quench different fluorophores with varying emission spectra, demonstrating the utility of multispectral detection with the system. Importantly, while the addition of the tetrazine-PEG-arm created a small bathochromic shift in the excitation peak by 6 and 3 nm in both dyes (λ_abs_ 496, λ_abs_ 593 nm) (**Fig. S1b**), the emission peak remained largely unaltered at λ_em_ 515 nm and λ_em_ 613 nm respectively (**Fig. 1c**).

Next, we evaluated the system’s performance by directly detecting protein-protein interactions in fixed cells. Our choice of target was the oncogene transcription factor complex cMyc/MAX (myc-associated factor X, MAX), selected due to its previous utility in evaluating and introducing the enzymatic proximity ligation assay to a wider audience (3). Furthermore, the unambiguous subcellular localization of the cMyc/MAX complex, primarily within the nucleus, as opposed to each individual protein, which is localized in both the nucleus and cytosol (**Fig. S2**), makes it an ideal candidate for assessing the specificity of our method.

Since rabbit-raised antibodies are easily accessible for diverse set of targets, we aimed to assess the feasibility of using two rabbit-raised antibodies in a single assay. Using antibodies from separate species could be limiting due to the comparatively smaller pool of available antibodies for non-rabbit species, potentially restricting the range of interactions that can be tested. Rabbit-raised primary antibodies for cMyc and MAX were targeted by secondary antibodies conjugated to the dienophile trans-cyclooctene (TCO, **Fig. S1e-g**). After washing, a quick incubation with the pegylated Y-shaped fluorogenic tetrazine probe generated a fluorescent signal restricted to the nucleus (**Fig. 1d**). This strong fluorogenic turn-on was only observed when both antibodies were present (**Fig. 1d-h,** *F*1,266 = 22.8,*P* < 0.0001). This is in stark contrast to ordinary immunofluorescence where combining antibodies did not generate an additative signal (,*F*_1,302_=3.18 *P* > 0.05). As expected the cMyc/MAX complex is localized in the nucleus because the heterodimer binds to DNA and acts as a transcription factor. In contrast, individual cMyc and MAX proteins, as visualized by standard immunofluorescence, can be found in both the nucleus and cytosol (**Fig. 1h, Fig. S2b-d**).

## FluoroProx is Antigen- and Interaction-Specific

While it is promising that primary antibodies from the same species can be used, this approach presents the potential for non-specific interactions and cross-reactivity. There is a possibility of these antibodies binding to each other due to their shared, similar species-specific epitopes, instead of binding to their intended targets. Such interactions can lead to false-positive results and inaccuracies in data interpretation. To enhance the evaluation of method specificity, we replaced the MAX antibody with a rabbit polyclonal IgG isotype control (**Fig. 2**). We quantified the fluorescent intensity in the nucleus as well as in the cytosol and samples where both cMyc and MAX primary antibodies were used conjoint in staining (*M* = 65.77, *SD* = 5.01) had twice the fluorescent intensity (*t* = 7.63, *P* < 3 × 10^−6^) compared to samples that were stained by cMyc and isotype control (**Fig. 2b**, *M* = 53.19, *SD* = 1.86) or isotype control alone (**Fig. 2c**, *t* = 6.79, *P* <1.2 × 10 ^−5^). Importantly, isotype control alone (*M* = 54.69, *SD* = 1.93) did not differ significantly from cMyc together with isotype control (*t* =-1.67, *P* = 0.11), indicating that cross-reactivity and unspecific binding is less of a concern.

**Figure 2.**
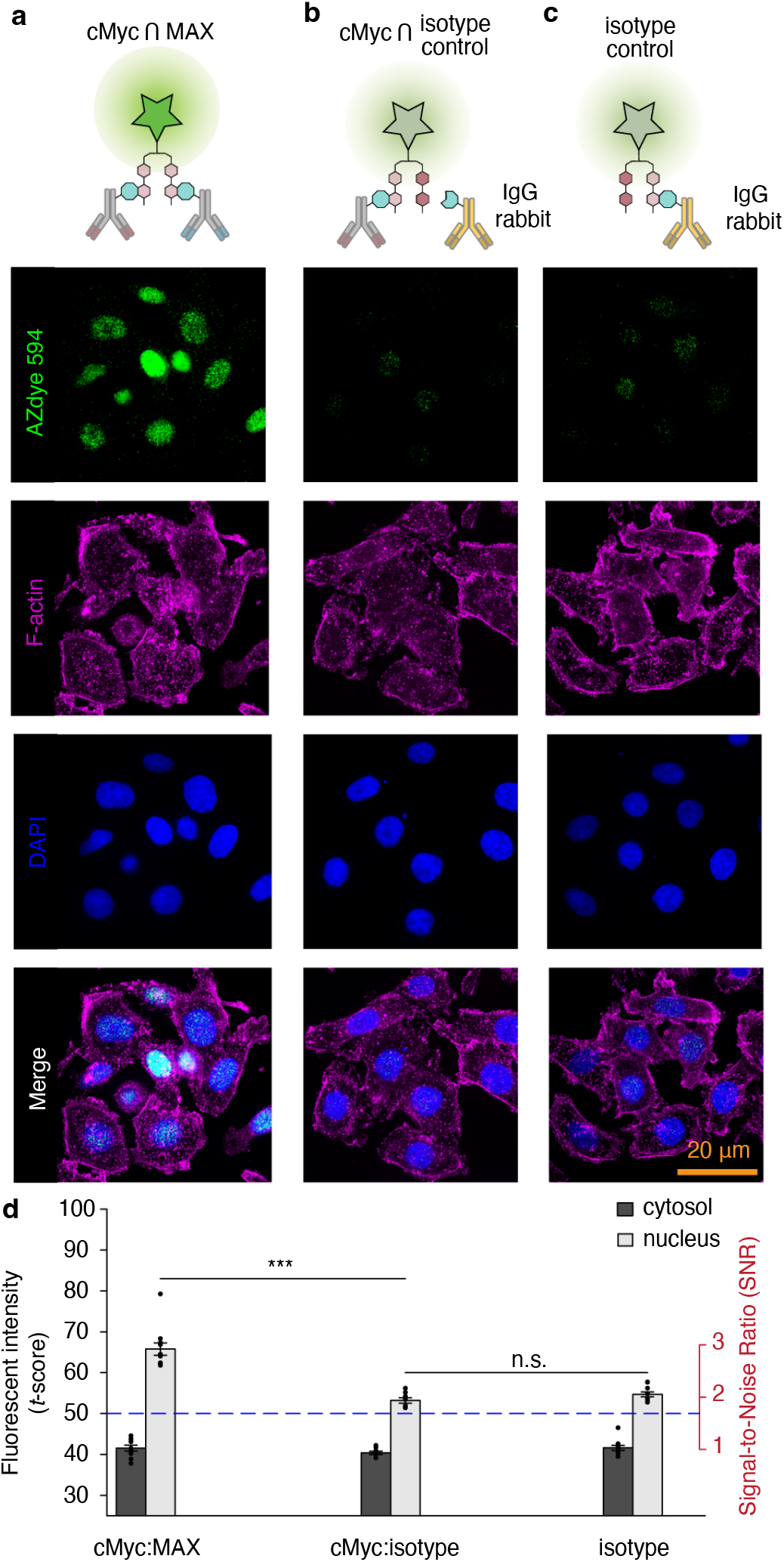
FluoroProx Signal is Antigen-Specific. **a)** Detection of cMyc:Max rabbit IgG primary antibodies. **b)** Negative control with same immunoglobulin isotype, rabbit IgG, as Max and cMyc. No significant interaction with cMyc is detected. **c)** Isotype control, rabbit IgG, alone exhibits same pattern as cMyc:isotype control. Faint granulated signal in the nucleus. Scale bar: 20 μm. **d)** Quantification of average fluorescent signal intensity in cytosol, dark gray, and nucleus, light gray.

Having observed that our method specifically detects protein-protein interactions subcellularly localized to the cell nucleus we decided to test the method on detection a protein-protein interaction pair localized to a different subcellular compartment. The membraine domain pair E-cadherin and β-catenin is a highly studied interactions due to its role in epithelial-mesenchymal transition. We targeted each protein using monoclonal primary antibodies with epitopes close to the site of interaction (**Fig. 3a**). A membrane localized fluorescent turn-on signal (*M* = 62.52, *SD* = 7.34) was only observed when both antibodies were used (**Fig. 3e**) whereas using anti-E-cadherin (*t* = 24.72, *P* < 0.0001; *M* = 50.91, *SD* = 2.62) or anti-β-cadherin alone (*t* = 24.47, *P* < 0.001; *M* = 54.75, *SD* = 2.87) yielded no signal boost.

**Figure 3.**
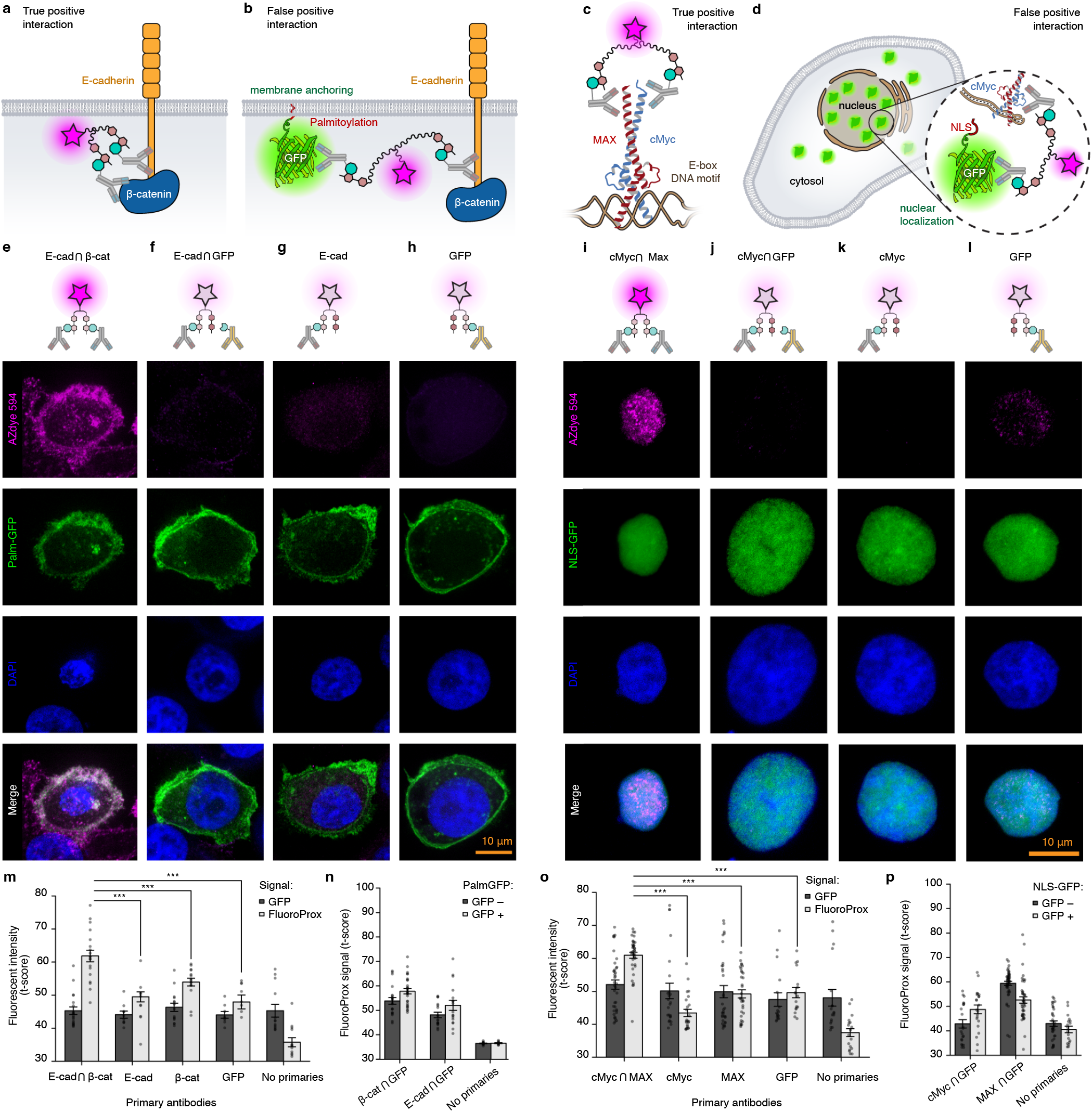
FluoroProx Probes Detect Interactions rather than Proximity. **a)** Targeting of the E-cadherin/β-catenin protein interaction localized to the cell membrane by monoclonal antibodies and FluoroProx probes. **b)** Schematic showing possible false positive interaction displayed by tethering GFP to the plasma membrane with a palmitoylation signal and targeting both E-cadherin and GFP with antibodies. **c)** Targeting cMyc/MAX protein interaction localized to the nucleus. **d)** Possible false positive interaction by localizing GFP to the nucleus with a nuclear localization signal (NLS). **e)** E-cad/β-cat interaction in Palm-GFP positive cell visualized by anti-E-cad and anti-β-cat antibodies. **f)** anti-E-cad and anti-GFP antibodies in Palm-GFP positive cell does not generate a strong fluorescent turn-on. **g)** Absence of fluorescent turn-on signal when only anti-E-cad antibodies are utilized.**h)** No fluorescent turn-on signal observed when only anti-GFP antibodies are employed. **i)** Visualization of the cMyc/MAX interaction in NLS-GFP positive cells using anti-cMyc and anti-MAX antibodies. **j)** Application of anti-cMyc and anti-GFP antibodies in NLS-GFP positive cells does not result in a significant fluorescent turn-on. **k)** Lack of a fluorescent turn-on signal when only anti-cMyc antibodies are applied. **l)** Absence of a fluorescent turn-on signal observed when only anti-GFP antibodies are used. **m)** Quantification of average nuclear fluorescent intensity in GFP (dark gray) and FluoroProx channel (light gray). **n)** No significant interaction between GFP and β-caterin or E-cadherin is obtained comparing PalmGFP-positive (GFP+, light gray) and PalmGFP-negative cells (GFP-, dark gray). **o)** Similar quantification as in m) but for NLS-GFP cells. GFP signal intensity (dark gray) and FluoroProx signal intensity (light gray). **p)** No significant signal boost when GFP antibody is used comparing NLS-GFP-positive (light gray) and negative cells (dark gray). Scale bars: 10 µm, Error bars: +/-one standard error of measurement.

Since we now had a pair of differentially localized protein-protein interactions in cMyc/MAX (nucleus) and E-cad/β-cat (membrane) we decided to use these two compartment localizations to evaluate our methods robustness towards spurious proximity-based interactions rather than true protein-protein interaction. By introducing a green fluorescent protein (GFP) tagged with N-terminal localization signals (palmitoylation signal from GAP43 or NLS from SV40) we could either localize GFP to the vicinity of E-cad/β-cat at the lipidmembrane (**Fig. 3b**) or near cMyc/MAX in the nucleus (**Fig. 3d**).

Cells expressing Palmitoylation-GFP (Palm-GFP) were analyzed by segmenting the membrane from the GFP channel, followed by measuring the average FluoroProx signal within the segmented membrane masks (**Fig. S4b**). Only in E-cad/β-cat antibody targeted cells did we observe a characteristic signal boost indicating protein-protein interaction (**Fig. 3e,m**) but not in cells targeted with anti-E-cad and anti-GFP or anti-β-catenin and anti-GFP antibodies (*F*_2,103_ = 1.12, *P* > 0.05; **Fig. 3f-h,m**).

Next, we examined if the introduction of GFP localized to the nucleus through a NLS element would induce any proximity-induced signals to cMyc. We did not observe any significant signal boost in NLS-GFP positive cells (**Fig. 3k-l**). These findings underscore the specificity and applicability of our method in discerning true protein-protein interactions within distinct subcellular compartments.

To determine whether our probes could detect a true protein-protein interaction, we fused GFP directly to E-cadherin and tested if this would be recognized as an interaction. In previous controls, GFP was merely localized to the same compartment as the protein of interest, which did not yield a positive signal. If FluoroProx indeed measures protein-protein interactions, this ‘synthetic protein-protein interaction’ should yield a significant boost in signal. This is also what we observe as indicated by a significant signal boost when both antibodies were present (*M* = 63.66, *SD* = 11.27) as opposed to either anti-E-cadherin (*t* =-6.3, *P* < 0.0001, *M* = 50.27, *SD* = 4.36) or anti-GFP alone (*t* = -2.96, *P* < 0.005, *M* = 53.68, *SD* = 13.81) (**Fig. 4**).

**Figure 4.**
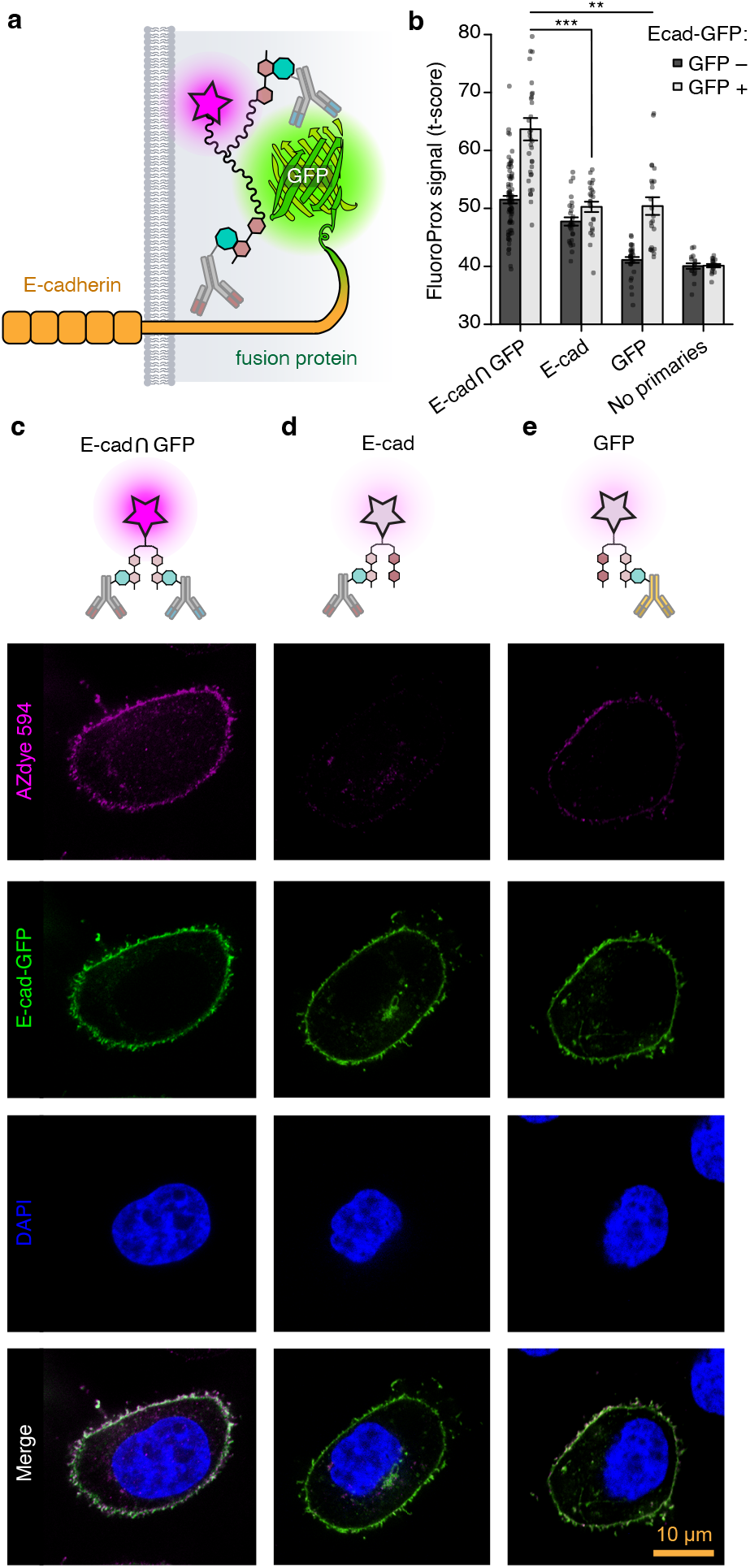
Detecting Fusion Proteins. **a)** E-cadherin was fused to GFP at the C-terminal and antibodies targeted for both GFP and E-cadhering was used. **b)** FluoroProx signal in GFP positive (light gray) or GFP negative (dark gray) cells when either anti-Ecad and anti-GFP or either antibody alone is used. *n* = 252 cells. **c)** anti-E-cadherin and anti-GFP antibodies. **d)** anti-E-cadherin antibody alone. **e)** anti-GFP antibody alone. FluoroProx594 (magenta), E-cadherin-GFP (green), counter stained with DAPI (blue). Scale bar: 10 µm.

Taken as a whole these findings highlight our method’s ability to distinguish true protein-protein interactions from proximity-induced events within distinct subcellular compartments. Notably, this discrimination is difficult to achieve using established in situ methods (17).

## Live Imaging using Genetic Code Expansion

For live imaging, we aimed to site-specifically label a protein capable of forming homodimers. OmoMYC is a synthetic protein developed to target and inhibit the activity of cMyc and its interaction with MAX (18). OmoMYC works by binding to cMyc and preventing its regulation of transcription and cell proliferation, which are involved in promoting cancerous growth. This inhibition of cMyc activity has shown promise in studies as a potential therapeutic approach for cancer treatment. When OmoMYC is present, it forms heterodimers with cMyc, thereby sequestering cMyc away from DNA. Additionally, it creates transcriptionally inactive homodimers and heterodimers with MAX, which occupy E-boxes (**Fig. 5a**), leading to the inhibition of transcription of cMyc targets. Therefore, we decided to tag the OmoMYC/OmoMYC dimers by site-directed labeling through genetic code expansion (**Fig. 5a**). A tamoxifen-inducible OmoMYC plasmid (19) was labeled for genetic code expansion by site-directed mutagenesis of a glycine codon at position 91 into an amber codon (OmomycTAG_91_-ER, **Fig. 5b**). We verified that the OmoMYC synthetic protein indeed inhibited cMyc/MAX interaction as measured by FluoroProx antibody staining with cMyc/MAX as the target (**Fig. 5c,d**). cMyc/MAX interaction was not only heavily reduced in tamoxifen treated cells (*M* = 37.92, *SD* = 1.73) compared to non-treated cells (*M* = 54.36, *SD* = 7.93, *t* = -15.23, *P* < 0.0001), but the few cMyc/MAX complexes that still existed were mainly localized in nucleolus or outside of the nucleus (**Fig. 5d**). Co-transfection of the OmomycTAG_91_-ER plasmid with a plasmid expressing a tRNA and its cognate aminoacyl-tRNA-synthetase pair (20, 21) together with the non-canonical amino acid (ncAA) azido-phenylalanine (azido-Phe or AzF) enabled site-directed labeling and strong nuclear localization when counter-stained with both a DNA and lipid membrane stain (**Fig. 5e-g**). We tracked a total of 24 individually labeled cells (*n* = 7 biological replicates) for on average forty minutes at one frame for every ten seconds. As expected from a homodimer binding to DNA E-boxes signal was restricted to the nucleus (**Fig. 5e**). Comparing the two FluoroProx dyes 488 and 594 in living cells we observed that 488 was more prone to photobleaching but both dyes produced qualatively similar labeling signal (**Fig. S4**). In conclusion, our study serves as a proof-of-concept for directly visualizing protein-protein interactions in living cells through the site-specific labeling of OmoMYC/OmoMYC dimers using genetic code expansion.

**Figure 5.**
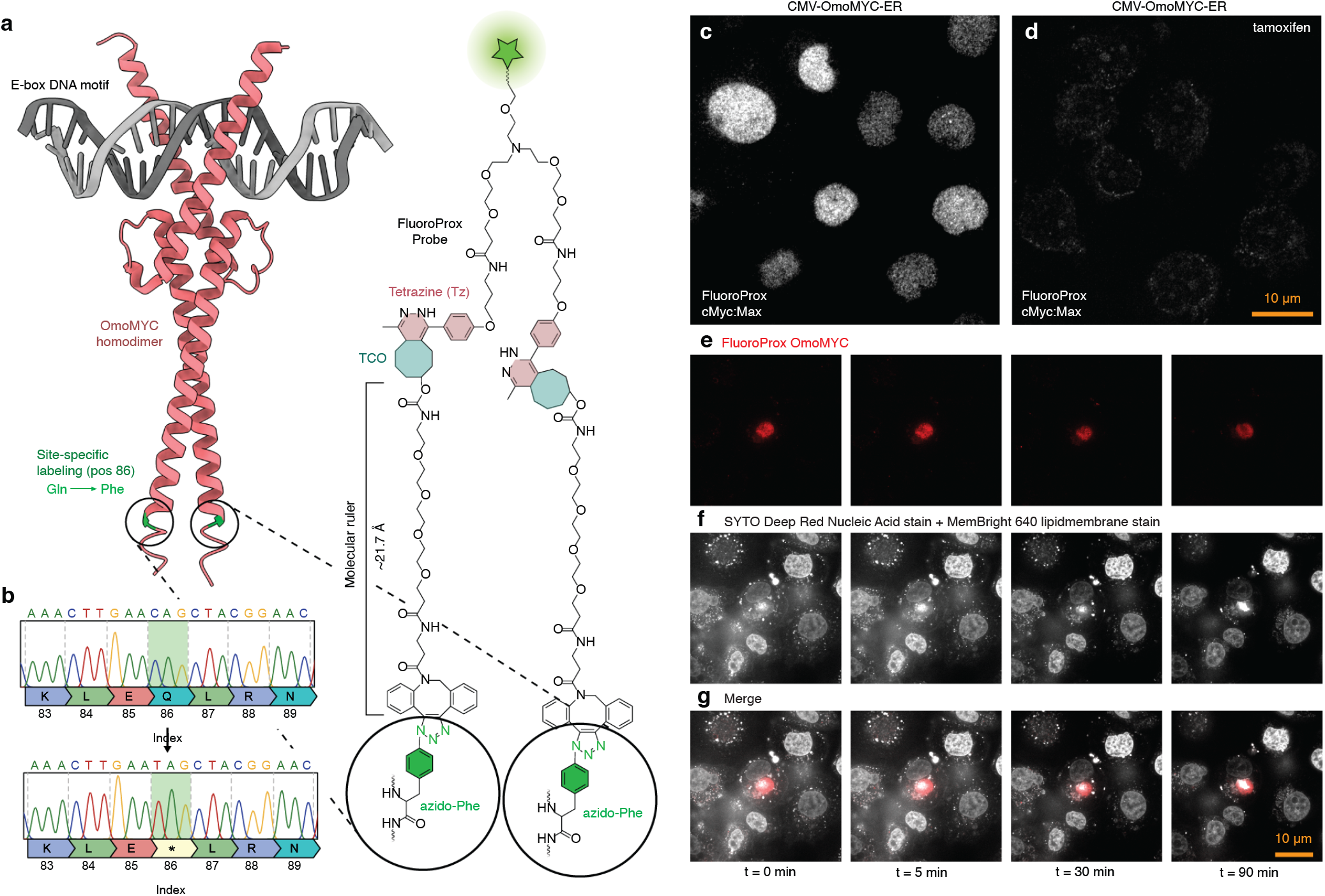
Live-cell Imaging of Protein-Protein Interactions. **a)** Representation of the crystal structure of the OmoMYC/OmoMYC dimer (PDB ID: 5I50) in red, bound to consensus E-box DNA in gray, with sites for site-directed mutagenesis highlighted in green. **b)** Sanger sequencing results for site-directed mutagenesis of amino acid 91 into an amber codon for genetic code expansion. **c)** OmoMYC-ER transfected PC3 cells that have not been tamoxifen-induced exhibit cMyc/MAX heterodimers visualized by FluoroProx probes. **d)** Tamoxifen induction of OmoMYC-ER not only inhibits cMyc/MAX dimers but also localizes any remaining weak signal to nucleoli or outside the nucleus. **e-f)** Live imaging of PC3 cells transfected with OmomycTAG_91_-ER sampled at 0.1 Hz. FluoroProx-594 imaged in red (**e**) and counterstained for DNA and lipidmembranes in far-red channel using SYTO Deep Red and MemBright 640 shown in gray (**f**). (**g**) Merged gray and red channel. Scale bar: 10 μm. A total of 24 OmoMYC positive cells were tracked individually on average for 38 min (*n* = 7 biological replicates).

## Discussion

The challenge of detecting protein-protein interactions within cells has long been a bottleneck in understanding cellular processes. Here we introduces a novel approach to tackle this challenge by employing fluorogenic probes with dual-tetrazine pegylated branched arms linked to a xanthene dye. The design of these probes addresses the limitations associated with existing methods, providing a promising avenue for compatibility in living cells without altering protein function via fluorescent tagging, while also being capable of detecting endogenous proteins where transgenic methods are not viable, such as clinical samples.

Our work highlights some specific limitations and suggests a clear path forward for developing this technology. First, the quenching process is incomplete. The spacer introduced by the bifurcated PEG-linker naturally keeps the tetrazines seperated from the fluorophore. Future research could explore different distances to find a better balance between steric hindrance and optimal physicochemical properties.

Another limitation is that probes that have only reacted with one moiety (mono-substrate) and not with both substrates will remain in a semi-quenched state, still detectable by microscopy, though dimmer. This signal could be fully quenched by washing with dabsyl-TCO quenchers. Additionally, the mono-substrate-bound probes could be visualized by allowing the free tetrazine arm to react with a fluorophore-conjugated TCO that emits at a third wavelength, such as far-red.

In addition to its current applications, the technology introduced in this study holds great promise for future advancements in the direct tracking of protein-protein interactions as well as protein-ligand interactions within living organisms. The dual-tetrazine pegylated branched arms linked to xanthene dye offer a unique opportunity to study protein-ligand interactions in real-time, providing a potential breakthrough in understanding drug kinetics and interactions directly *in vivo*. The non-invasive nature of the fluorescent probes mitigates concerns associated with altering protein function or cellular processes, making it an ideal candidate for investigating the dynamic interplay between drugs and their target proteins within the complex biological environment. This capability to observe and quantify protein-ligand interactions in living systems has profound implications for drug development, allowing researchers to gain insights into the temporal aspects of drug binding, dissociation, and overall pharmacokinetics. The presented technology, with its dual-substrate recognition mechanism, not only expands our toolbox for studying interactions but also opens up new avenues translating results direcly between experimental transgenic systems and endogenous detection in samples where transgenic manipulation is not feasible, such as clinical samples.

## Material & Methods

### Image analysis

Raw images were acquired as 16-bit z-stacks but analysis was done on maximum intensity projections in 16-bit raw tiff format. For cell segmentation using phalloidin staining we utilized a simple U-net with weights for cell boundaries (22). For segmentation of DAPI we used StarDist (23).

### Statistical analysis

All statistical computations were done with R the statistical programming language. Fluorescent intensity was normalized between replicates to a mean of 50 and standard deviation of 10 by t-score transformation:

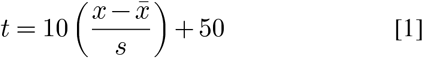

Where:

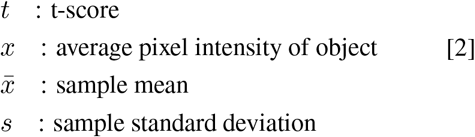

Statistical tests performed were either two-sided t-tests or ANOVA.

### Cell culture

Prostate cancer cells (PC3), osteosarcoma cells (U2-OS) and human embryonic kidney cells (HEK 293) were cultivated in Advanced Dulbecco’s Modified Eagle Medium (DMEM) (12491015, Thermo Fisher, Waltham, MA) supplemented with 10% (v/v) fetal bovine serum (10106-147, Thermo Fisher, Waltham, MA), 1% (v/v) Penicillin, Streptomycin and L-glutamine (PSG) (10378016, Thermo Fisher, Waltham, MA). The cells were incubated at 37°C in a humidity controlled environment under a 5% CO2 atmosphere.

### Fluorogenic probe synthesis

N-(Azido-PEG1)-N-bis(PEG2-Methyltetrazine-propylamine) (BP-28748, Broad Pharma Inc.) was dissolved in anhydrous DMSO (276855, Sigma-Aldrich Gmbh) to a stock concentration of 10 mM. Likewise, fluorophore of choice (AZdye488-DBCO or AZdye594-DBCO, 1278-1 or 1298-1 Click Chemistry Tools Inc.) was dissolved in DMSO to a a stock concentration of 10 mM. Initially the pegylated bis-tetrazine arms were conjugated to fluorophore of choice through azide/DBCO Strain-promoted Azide-Alkyne Click (SPAAC) reaction in DMSO at 1:1 molar equivalents for 24h in room temperature. Conjugation was verified by fluorescent quenching of the fluorophore by the tetrazines as well as liquid chromatography–mass spectrometry (LC–MS).

### Antibody conjugation

Antibody conjugation was performed using TFP-esters according to the following protocol. First, antibody purification was carried out to remove interfering compounds if present such as BSA, azide, or glycine often added by the manufacturer. This involved washing the antibody in 1× PBS using Amicon ultra 30K 0.5mL columns (UFC5030, Merck GmbH), followed by collection of the purified antibody and determination of its concentration using a NanoDrop spectrophotometer at 280 nm.

For ester preparation, TFP-ester (TCO-PEG4-TFP, BP-40298, Broad Pharma Inc.) was dissolved in anhydrous DMSO to a concentration of 10 mg/mL. The antibody was then labeled at 2 mg/mL in 0.5 M carbonate/bicarbonate buffer (pH 8.75) at various molar ratios of ester to antibody (3×, 9×, 15×). After incubation at room temperature for 1h in the dark, unreacted ester was quenched by the addition of 1M Tris buffer (pH 8.0) to a final concentration of 100 mM and incubated on ice for 15 min.

The labeled antibody conjugates were purified and concentrated using Zeba Spin Desalting Columns 40K MWCO (11796436, ThermoFisher Inc.) and Amicon ultra 30K 0.5mL centrifugation column (UFC5030, Merck GmbH). Excess ester was removed through multiple wash steps with 1× PBS, and the final conjugates were collected and their concentrations and degree of labeling was determined by NanoDrop and SDS-PAGE analysis. The resulting antibody conjugates were diluted to a final concentration of 2 mg/mL with 1× PBS and stored at 4 degrees Celsius for subsequent experiments. Full protocol and all reagents is accessible here: https://www.furthlab.xyz/antibody_conjugation.

### Fluorescent spectrophotometry

FluoroProx probes at a stock concentration of 5 mM were diluted in 1xPBS pH 7.5 to a concentration of 2 uM for a total volume of 2.5 mL in polystyrene disposable cuvettes with four clear faces (634-8530, VWR) and emission was measured on a Fluorolog SPEX TCSPC Horiba fluorescence spectrophotometer (ex. wavelength 488 and 594 nm, 0.1 sec. integration time, 1 nm slit). Absorbance was measured by UV-Vis spectrophotometry on a Denovix DS-11 spectrophotometer.

### Liquid chromatography–mass spectrometry (LC–MS)

LC-MS analysis was carried out using electrospray ionization (ESI) and a C18 column (50×3.0 mm, 2.6 μm particle size, 100 Å pore size) with acetonitrile/water in 0.05% aqueous formic acid as mobile phase.

***MS(ESI)***, m/z calc’d for C_81_H_92_N_18_O_20_S_2_: 1696.59; found 1697.7 [M+H]^+^, 849.0 [M+2H]^2+^.

***MS(ESI)***, m/z calc’d for C_95_H_107_N_18_O_20_S_2_: 1884.74; found 1885.7 [M+H]^+^, 943.1 [M+2H]^2+^.

FluoroProx488 *“AZDye488-DIBOT-PEG1-N-bis(PEG2-Tz)”*

**6-amino-9-(2-carboxy-4-((3-(1-(19-(4-(6-methyl-1,2,4,5-tetrazin-3-yl)phenoxy)-6-(2-(2-(3-((3-(4-(6-methyl-1,2,4,5-tetrazin-3-yl)phenoxy)propyl)amino)-3-oxopropoxy)ethoxy)ethyl)-15-oxo-3,9,12-**

**trioxa-6,16-diazanonadecyl)-1,9-dihydro-8H-dibenzo[b,f][1,2,3]triazolo[4,5-d]azocin-8-yl)-3-oxopropyl)carbamoyl)phenyl)-3-iminio-5-sulfo-3H-xanthene-4-sulfonate**

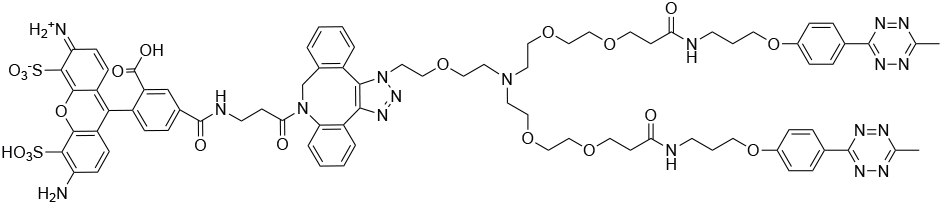

FluoroProx594 *“AZDye594-DIBOT-PEG1-N-bis(PEG2-Tz)”*

**(6-(2-carboxy-4-((3-(1-(19-(4-(6-methyl-1,2,4,5-tetrazin-**

**3-yl)phenoxy)-6-(2-(2-(3-((3-(4-(6-methyl-1,2,4,5-tetrazin-3-yl)phenoxy)propyl)amino)-3-oxopropoxy)ethoxy)ethyl)-15-oxo-3,9,12-trioxa-6,16-diazanonadecyl)-1,9-dihydro-8H-dibenzo[b,f][1,2,3]triazolo[4,5-d]azocin-8-yl)-3-oxopropyl)carbamoyl)phenyl)-1,2,2,10,10,11-hexamethyl-10,11-dihydro-2H-pyrano[3,2-g:5,6-g’]diquinoline-1-ium-4,8-diyl)dimethanesulfonate**

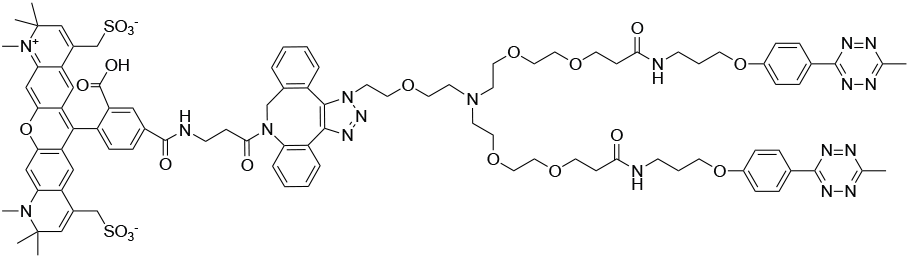

**Immunofluorescence**. PC3 cells (7.5 × 10 ^3^) were seeded on 35 mm glass-bottom dishes (81218, Ibidi). Once the cells reached approximately 70% confluence, the growth medium was replaced with 1× PBS (AM9625, Thermo Fisher Scientific), and cells were washed three times with ice-cold PBS. Cells were then fixed with 4% (v/v) formaldehyde (PFA; 28908, Thermo Fisher Scientific) in 1×PBS for 15 minutes. Following fixation, cells were washed three times with PBST containing 0.25% (v/v) Triton-X100 (T8787, Sigma-Aldrich) in 1×PBS and permeabilized at room temperature for 10 minutes. To remove detergent, cells were washed three additional times with 1× PBS.

Blocking was performed using 1% (v/v) goat serum (GOA-1A, Capricorn Scientific GmbH) diluted in 1× PBS containing 0.1% (v/v) Tween-20 (P9416, Sigma-Aldrich) for 1 hour at room temperature. After blocking, cells were washed three times with PBST (0.1% v/v Tween-20 in 1× PBS), with each wash involving a 5-minute incubation.

Primary antibodies, prepared as detailed in Table 1, were diluted in a solution containing 1% (w/v) BSA (A9647, Sigma-Aldrich) in 1× PBS with 0.1% (v/v) Tween-20. The cells were incubated with the primary antibodies overnight at 4^∘^C on a plate shaker.

**Table 1.** List of antibodies.

| Antibody | Vendor | Catalog Number | LOT Number | Dilution | Host | Isotype | Clonality |
| --- | --- | --- | --- | --- | --- | --- | --- |
| Alexa488 Chicken Anti-Rabbit IgG (H+L) | Invitrogen | A-21441 | 2549869 | 1:1000 | Chicken | IgY | Poly |
| Alexa647 Goat Anti-Mouse IgG (H+L) | Invitrogen | A-21235 | LOT456 | 1:500 | Goat | IgG | Poly |
| Goat Anti-Rabbit IgG(H+L) | Invitrogen | 31210 | 086151L | 1:1000 | Goat | IgG | Poly |
| Goat anti-Mouse IgG (H+L) | Invitrogen | A16080 | 95-87-012723 | 1:1000 | Goat | IgG | Poly |
| Goat anti-Rabbit IgG (H+L) | Invitrogen | 31212 | YJ4089381 | 1:1000 | Goat | IgG | Poly |
| Rabbit IgG Isotype Control | Invitrogen | 10500C | XJ358702 | 1:12.5k | Rabbit | IgG | Poly |
| GFP tag Polyclonal antibody | Proteintech | 66379-1-ig | 10011552 | 1:100k | Rabbit | IgG | Poly |
| c-Myc Recombinant Rabbit mAb (27H46L35) | Invitrogen | 700648 | 2561984 | 1:500 | Rabbit | IgG | Mono |
| c-Myc Mouse Monoclonal IgG (9E10) | Santa Cruz | 9E10 | C2823 | 1:300 | Mouse | IgG | Mono |
| Rabbit Anti-MAX Antibody | AtlasAntibodies | HPA003474 | A115844 | 1:500 | Rabbit | IgG | Poly |
| MAX Recombinant Rabbit Monoclonal (103) | Invitrogen | MA5-29412 | YL4165092 | 1:20k | Rabbit | IgG | Mono |
| B-catenin(D10A8) XP - Rabbit mAb | Cell Sign. Tech. | AD10A8 | 9 | 1:1500 | Rabbit | IgG | Mono |
| B-catenin(E-5) Mouse mAb | Santa Cruz | SC-7963 | JO622 | 1:500 | Mouse | IgG | Mono |
| E-cadherin(24E10) Rabbit mAb | Cell Sign. Tech. | 24E10 | 15 | 1:500 | Rabbit | IgG | Mono |

**Table 2.** List of plasmids.

| Plasmid | Addgene ID | Description | Provided by | Reference |
| --- | --- | --- | --- | --- |
| CAG-NLS-GFP | 104061 | GFP N-terminal NLS from SV40 | Gradinaru | (24) |
| pCAG-mGFP | 14757 | GFP N-terminal palmitoylation signal from GAP43 | Cepko | (25) |
| CAAX-EGFP | 86056 | Farnesylation motif of HRAS with N-terminus EGFP | Lu | (26) |
| E-cadherin-GFP | 28009 | GFP fused to E-cadherin C-terminal intracellular domain | Stow | (27) |
| pAS_4xBstTyrT(CUA)_EcoTyrRS-FLAG | 140018 | E.coli tRNA cassette | Elsässer | (20) |
| pCS Omomyc-MER | 113170 | OmoMYC with N-terminal ER for tamoxifen induction | Nasi | (18) |
| pCbs FlagOmomyc | 113168 | OmoMYC with N-terminal FLAG-tag | Nasi | (19) |

The following day, cells were washed three times with PBST, with each wash lasting 5 minutes at room temperature. Secondary antibodies (Alexa Fluor 488 Chicken Anti-Rabbit IgG (H+L); Lot: 2549869, A21441, Thermo Fisher Scientific) were diluted 1:1000 in PBST with 1% (w/v) BSA and incubated for 1 hour at room temperature. Cells were then washed again in PBST using the same protocol as above, followed by a thorough wash in 1× PBS.

### Fluorogenic Dual-Substrate Recognition Assay in Fixed Cells

Confluent cells were fixated by incubating the samples with 4% formaldehyde (PFA) for 15 minutes. Following fixation, the cells were washed three times with PBST (PBS containing 0.1% Tween-20). Permeabilization was carried out by incubating the cells in 0.25% Triton-X in PBS for 15 minutes, followed by an additional three washes with PBST. To block nonspecific binding sites, the cells were incubated in 10% serum solution for 1 hour, followed by three washes with PBST. Samples were then incubated with primary antibodies overnight at 4°C. The cells were subsequently washed three times with PBST before a 1-hour incubation at room temperature with TCO-conjugated (1398-2, Click Chemistry Tools Inc.) secondary antibodies (31210, 31212, or A16080 ThermoFisher). After washing three times with PBST, FluoroProx probes were added at a final concentration of 200 nM in 1 × PBS for 20 minutes. Following a final wash step with PBST, the samples were prepared for imaging.

### Image acquisition and processing

Cells were imaged on a Nikon TE2 equipped with a CrestOptics XLight V3 spinning disc confocal. A fluorescent light source was provided by Lumencor’s CELESTA Quattro Light Engines with an arrays of 5 individually addressable solid-state lasers (Lumencor CELESTA Quattro nIR 5ch with Despeckler, 405/12, 476/12, 545/12, 637/12 and 748/12) and images were acquired with a Kinetix sCMOS Camera (Photometrics) through a oil immersion 100x objective. The spinning disc was equipped with a dichroic beam splitter (FF421/491/567/659/776-Di01, Semrock Inc.) together with a penta-band bandpass filter, emitter (FF01-391/477/549/639/741, Semrock Inc.) and exciter (FF01-391/477/549/639/741, Semrock Inc.), optimized for DAPI/FITC/TRITC/Cy5/Cy7 Lumencor CELESTA. Single-band bandpass filters included (FF02-438/24; FF01-511/20; FF01-595/31; FF02-685/40; FF01-819/44). A complete description of the setup can be accessed from FPbase.org:https://www.fpbase.org/spectra/?s=6645,6642,6646,6644,780,884,1090,1052,987,4707,4708,4709,

### Plasmids

Translocation of fluorophores to the nucleus was achieved by tagging the N-terminal of GFP with Simian Vacuolating Virus 40 (SV40) Nuclear Localisation Signal, CAG-NLS-GFP (24) (Addgene plasmid #104061) was a gift from Viviana Gradinaru (California Institute of Technology). The cell membrane reporter used was N-terminal tagged GFP with Palmitoylated Growth Associated protein 43 (GAP-43), pCAG-mGFP (25) (Addgene plasmid #14757) was a gift from Connie Cepko (Harvard Medical School). Additionally, farnesylated HRAS-EGFP (26) fusion protein (CAAX-EGFP, Addgene plasmid # 86056) provided dynamic visualization of the Golgi and plasma membrane during imaging, this construct was kindly provided by Lei Lu (Nanyang Technological University).Interactions between β-Catenin and E-cadherin was studied through E-cadherin-GFP (27) (Addgene plasmid # 28009) was a gift from Jennifer Stow (University of Queensland). Aminoacyl-tRNA synthetase and non-sense tRNA for incorporation of p-azido-phenylalanine was translated from pIRE4-Azi (21) (Addgene plasmid # 105829) and pAS_4xBstTyrT(CUA)_EcoTyrRS-FLAG (20) (Addgene plasmid # 140018) were gifts from Irene Coin (Leipzig University) and Simon Elsässer (Karolinska Institute) respectively. The Tamoxifen-inducible fusion-protein of Omomyc, pCS Omomyc-Mer (Addgene plasmid # 113170) was a gift from Sergio Nasi (CNR-IBPM). A full list of plasmids can be seen in Table .

### Plasmid Purification

Plasmid-carrying bacterial cells were transferred from their respective agar stabs onto agar plates containing 100 µg/mL of Ampicillin or 50µg/mL of Kanamycin and incubated overnight at 37°C (DH5alpha) and 27°C (NEB stable). Alternatively, purified plasmids were amplified through transformation into electrocompetent E.coli (DH10B) through electroporation using Amaxa® Nucleofector II and subsequently seeded on solid agar containing the appropriate antibiotics and incubated at 37°C. Colonies of bacteria were subsequently harvested and cultured in 5 or 25 ml of liquid LB medium overnight. For smaller preparations, the tubes containing 5 mL of the cultured cells were centrifuged at max RPM for 1 min after which, the supernatant was removed and the pellet resuspended in in an aqueous buffer of 50 mM Tris HCl (ph 8), 10mM EDTA and 100 µg/ml RNase A. Lysis of bacterial cells was achieved using an alkaline solution containing 200 mM NaOH with 1% SDS (v/w) diluted in nuclease-free water. After incubation at RT for 5 min, 3M Sodium Acetate was added prior to centrifugation at max RPM for 10 min. The supernatant was transferred to new tubes and the plasmid DNA was precipitated using cold 2-propanol on ice for 5 min before being centrifuged at 4000 rcf for 5 min. Having discarded the supernatant, the pellet was washed with 70% ethanol before undergoing another centrifugation at 4000 rcf for 5 min. After the supernatant was discarded and the pellet dried from any residual ethanol, the plasmid DNA (pDNA) was resuspended in nuclease-free water.

Purification of 25 mL bacterial cultures was performed using PureLink HiPure Plasmid Midiprep Kit (K210004, Thermo Fisher, Waltham, MA) according to the manufacturer’s instructions. The results of the pDNA-extraction was subsequently confirmed by Gel-Electrophoresis on 1% Agarose gel, stained with 1 µl/mL SYBR™ Safe DNA Gel Stain (10,000X) (S33102, Thermo Fisher, Waltham, MA).

### Electroporation

PC3, U2-OS and HEK 293 cells were seeded and incubated overnight in Advanced DMEM containing 10% FBS, 100 IU/mL Penicillin, 10 µg/mL Streptomycin and 2mM L-Glutamate (Life Technologies, Bleiswijk, Netherlands #10378016) at 37°C in an atmosphere of 5% CO2. Upon reaching 70-80% confluency, cells were washed with 1xPBS before being detached by treatment with 5% Trypsin. After harvesting, the cells were concentrated using centrifugation and the pellet was subsequently resuspended in 1ml of culture medium. The transfection was carried out using Amaxa® Cell Line Nucleofector® Kit V. Approximately 106 cells (per reaction) were transferred to new 1.5ml tubes and centrifuged at max RPM for 10 min after which, the supernatant was removed completely. The cell-pellet was suspended in 100µl Cell Line Nucleofector® Solution V and 2µg of pDNA was added to the mixture. The entire sample was then transferred over to transfection cuvettes and placed in the cuvette holder of the machine (Amaxa® Nucleofector II). Electroporation was carried out using the cell appropriate program (T-013) after which, the solution was gently transferred from the cuvettes and plated onto 8-well chambers (CellVis, Mountain View, CA, Cat. #C8-1.5H-N). Full protocol can be accessed here: https://www.furthlab.xyz/lonza_nucleofection

### Lipotransfection

After reaching 70-80% confluence, PC3, U2-OS and HEK 293 cells were harvested and plated on 35 mm glass-bottom dishes (IBIDI, Gräfelfing, Germany, Cat. 81218-200) or 60 mm plastic dishes (Corning,) at a concentration of 2 × 10 ^4^ and 5 × 10 ^4^ cells/mL in full DMEM and incubated for 24h. The cells were thereafter washed 3 times with HBSS for 5 min before replacing the medium with Opti-Mem. Samples were subsequently transfected using either Lipofectamine or TransIt-293 at a ratio of 1:3 (DNA:transfectant).

### Polyethylenimine (PEI) transfection

A 1 µg/µL PEI solution was prepared by dissolving 100 mg of PEI powder in 90 mL of nuclease-free water (PEI MAX® -Transfection Grade Linear Polyethylenimine Hydrochloride (MW 40,000, 24765, Polysciences, Inc.). The pH was adjusted to 7.0 using 1 M HCl. The solution was stirred on a magnetic stirrer until the powder fully dissolved. The final volume was adjusted to 100 mL, and the solution was filtered through a 0.22 µm filter to remove any undissolved particles. The filtered solution was aliquoted and stored at -80°C.

Cells were grown to 70-80% confluence. For each transfection reaction, 1/20 of the total reaction volume was prepared with Opti-MEM. Plasmid DNA was added at 1 µg per mL of medium and mixed by vortexing. PEI stock solution was added at a 3:1 PEI ratio (w/w) and gently mixed. The mixture was incubated at room temperature for 10-15 minutes.

The transfection complex was added dropwise to the cells, ensuring even distribution. The vessel was gently rocked to disperse the particles. Optimization of the PEI ratio was performed using a 24-well plate with varying parameters to determine the optimal conditions for each cell line and plasmid batch.

### Genetic Code Expansion

Site-Directed Mutagenesis (SDM) was performed on the pCS Omomyc-Mer plasmid to introduce an amber codon on position 86 of the amino acid sequence of the insert. The primers used to modify the plasmid (Fwd: CTACGGAACTCTTGTGCGTAA; Ref: CTATTCAAGTTTGTGTTTCAACTG) were ordered from Integrated DNA Technologies (Leuven) at 25 nmole scale and reconstituted into nuclease-free water to a 100 µM stock concentration. The PCR was carried out using the Q5 Site-Directed Mutagenesis kit (E0554S, New England Biosciences, Ipswitch, MA) for 25 µl reaction containing 25 ng of template plasmid, 12.5 µl Q5 Hot Start High-Fidelity Master Mix (2x), 9 µl nuclesefree water and 1.25 µl 10 µM forward and reverse primers. The thermal cycle program included 25 cycles consisting of 98°C for 10 s, 61°C 30 s and elongation at 72°C for 1 min 45 s. Products were detected through gel electrophoresis on 1% agarose gel before re-circularizing the plasmid using KLD enzyme mix (M0554S, New England Biosciences, Ipswitch, MA). The modified plasmid was subsequently transformed into competent bacterial cells (E.coli) through electroporation and underwent selective cultivation on LB-plates containing 100 µg/mL Ampicilin. Single colonies were transferred to liquid LB for enrichment following pDNA isolation as described earlier. Samples of plasmid from mono-clonal origin were sent off for Sanger sequencing to validate the nucleotide substitution.

Harvesting of cells was performed as previously described and incubated in DMEM supplemented with 200 nM 4-azido phenylalanine (AzF) (CLK-AA001-10, Jena Bioscience, Jena, Germany) for 30 minutes prior to transfection. Co-transfection was performed according to protocol, with 1µg each of pIRE4-Azi (21) or pAS_4xBstTyrT(CUA)_EcoTyrRS-FLAG (20) and pCS Omomyc-ER (18) as a substitute for the 2µg of the respective plasmid used in the previous section. 4h post-transfection, the medium covering the seeded cells was removed by aspiration and replaced by 1µM AzF in DMEM and incubated for 48h.

### Live Imaging of GCE-modified cells

The medium containing ncAA was removed from the plates and the cells were washed with DMEM to remove any excess amino acids. TCO-PEG4-DBCO (BP-24160, Broad Pharma, San Diego, CA) reconstituted in DMSO to a 10 mM stock was further diluted in DMEM to a final concentration of 100 nM before being added to each sample and incubated for 45 min. The cells were then washed in DMEM three times over 45 min. FluoroProx probe (AZdye594-DIBOT-PEG1-N-bis(PEG2-Tz)) 50 nM in DMEM with 1X Probenecid (P36400, Thermo Fisher, Waltham, MA) was subsequently added to each well and incubated for 1h followed by another wash-step as described above. Counterstaining of the nucleus and membrane was performed 30 min before imaging the cells using 2µM SYTO™ Deep Red Nucleic Acid Stain (S34900, Thermo Fisher, Waltham, MA) and 0.1 µM MemBright 640 (SCT085, Sigma-Aldrich, Saint Louis, MO) in DMEM with 1X Probenecid respectively for 30 min. The samples were rinsed once with DPBS before replacing the DMEM with Leibowitz’s L-15 Medium (11415064, Thermo Fisher, Waltham, MA) containing 10% FBS, 1% PSG to allow for long-term microscopy without CO2 equilibration. Activation of the fused Estrogen-receptors was performed by addition of Tamoxifen to the cell medium drop wise to reach a final concentration of 0.5 µM.

## Acknowledgements

We gratefully acknowledge V. Agmo Hernandez for his expert assistance in helping us set up fluorescence spectrophotometry. We thank P. Wijethunga for providing advice and technical help. This work was supported by the a NARSAD Young Investigator Grant from the Brain & Behavior Research Foundation (29810), the Chan Zuckerberg Initiative (239942) and the Swedish Research Council (2022-02706), and SciLifeLab. Conflict of interest: D.F. serves on the scientific advisory board of Navinci Diagnostics AB.

---

**Figure S1.**
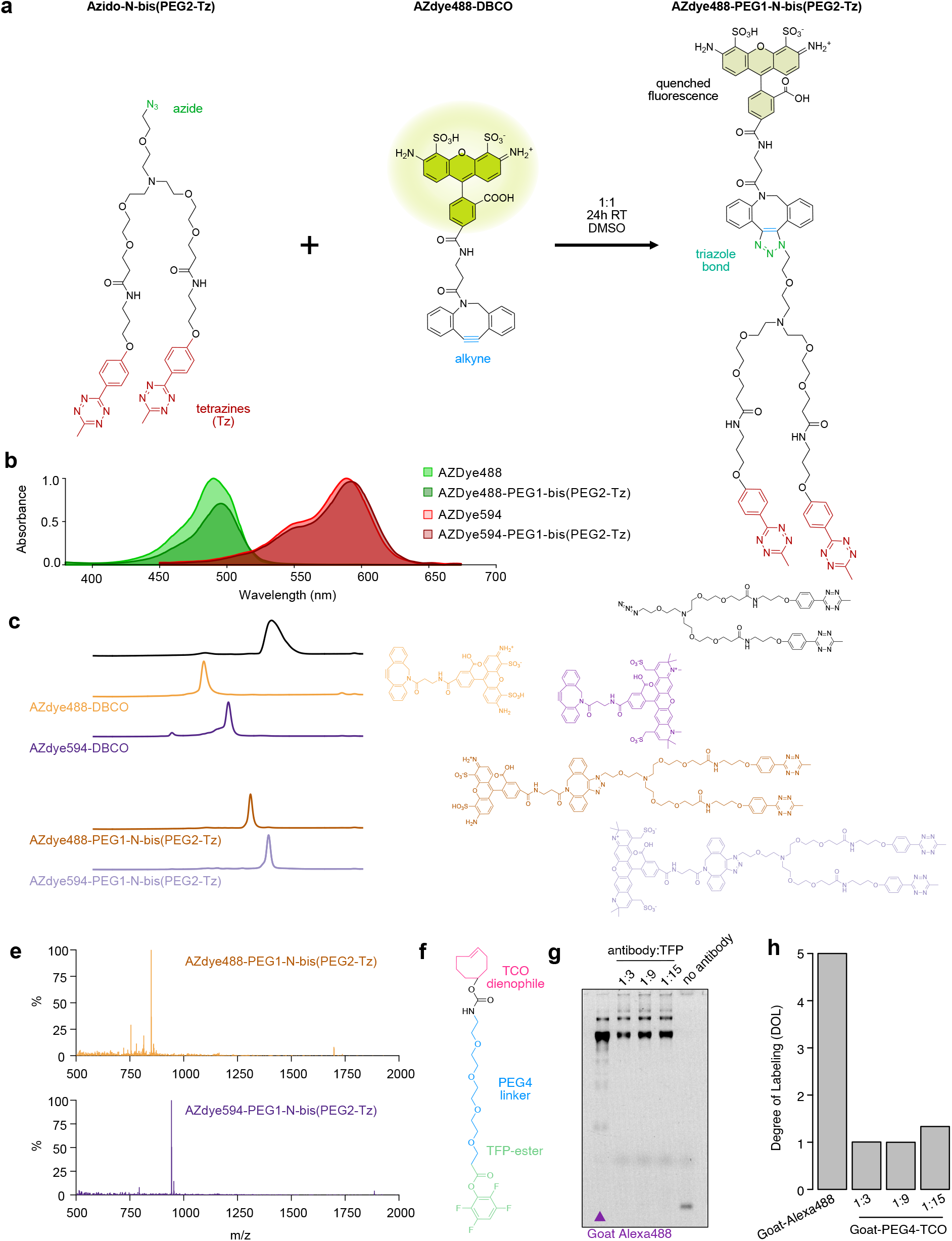
Synthesis of FluoroProx Probes. **a)** Strain-promoted azide-alkyne cycloaddition (SPAAC) between N-(Azido-PEG1)-N-bis(PEG2-Methyltetrazine-propylamine) (BP-28748, Broad Pharma Inc.) and DBCO (dibenzocyclooctyne) labeled fluorophore (1278, Click Chemistry Tools Inc.) performed in equimolar concentrations in anhydrous DMSO at room temperature overnight. **b)** Absorbance spectrum (excitation) of AZDye488-DBCO and AZDye594-DBCO before and after conjugation to N-(Azido-PEG1)-N-bis(PEG2-Methyltetrazine-propylamine). **c)** Liquid chromatography–mass spectrometry (LC–MS) chromatogram of reactants (AZDye488-DBCO, AZDye594, and N-(Azido-PEG1)-N-bis(PEG2-Methyltetrazine-propylamine)) and products (AZDye488-DIBOT-PEG1-N-bis(PEG2-Tz) and AZDye594-DIBOT-PEG1-N-bis(PEG2-Tz)). Structure of individual molecules shown to the right for clarity. **d)** Mass spectrogram of AZDye488-DIBOT-PEG1-N-bis(PEG2-Tz) and AZDye594-DIBOT-PEG1-N-bis(PEG2-Tz) from LC-MS in c). **e)** TCO-PEG4-TFP Ester used to label primary aliphatic amines on secondary antibodies. **f)** SDS-PAGE of TCO-conjugated antibodies and tetrazine-AZdye488 labeling. Lane 1 commercially conjugated goat secondary antibody. Lane 2: antibody to TCO-PEG4-TFP molar ratio of 1:3. Lane 3: molar ratio of 1:9. Lane 4: molar ratio of 1:15. Lane 5: tetrazine-AZdye488 alone. **g)** Degree of Labeling (DOL) of TCO-PEG4 conjugated antibodies.

**Figure S2.**
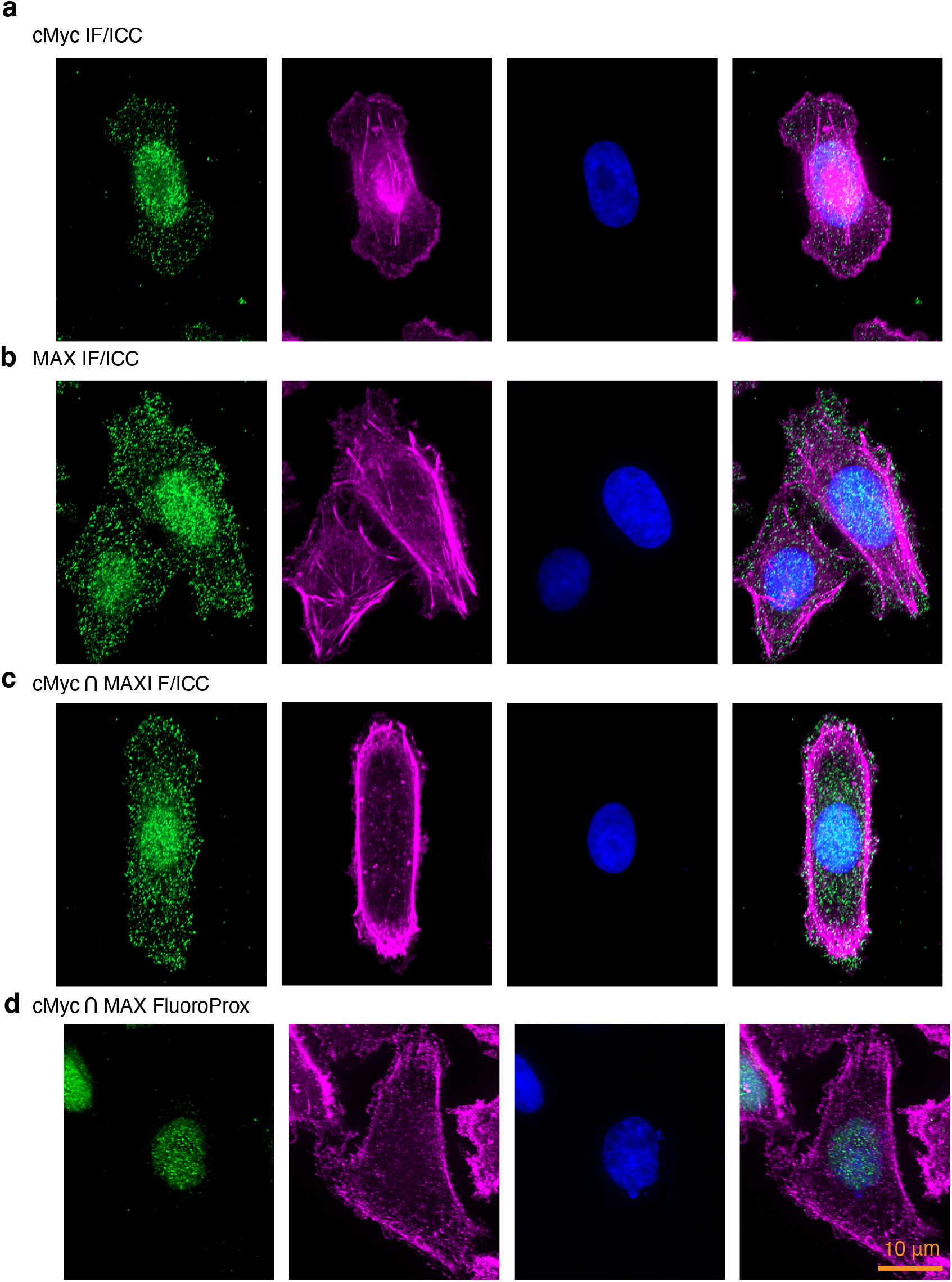
Comparison of IF/ICC to FluoroProx. **a)** Immunofluorescence of cMyc using monoclonal rabbit primary antibody visualized by polyclonal goat Alexa488-conjugated secondary antibody (green). Counter stained with DAPI (blue) and Phalloidin-iFluor594 (magenta). **b)** MAX immunofluorescence using polyclonal MAX antibody visualized by polyclonal goat Alexa488-conjugated secondary antibody (green). Counter stained with DAPI (blue) and Phalloidin-iFluor594 (magenta).**c)** cMyc and MAX immunofluorescence combined using monoclonal rabbit primary antibody for cMyc and polyclonal MAX antibody visualized by polyclonal goat Alexa488-conjugated secondary antibody (green). Counter stained with DAPI (blue) and Phalloidin-iFluor594 (magenta).**d)** cMyc and MAX FluoroProx combined using monoclonal rabbit primary antibody for cMyc and polyclonal MAX antibody visualized by FluoroProx594 (green). Counter stained with DAPI (blue) and Phalloidin-iFluor594 (magenta).

**Figure S3.**
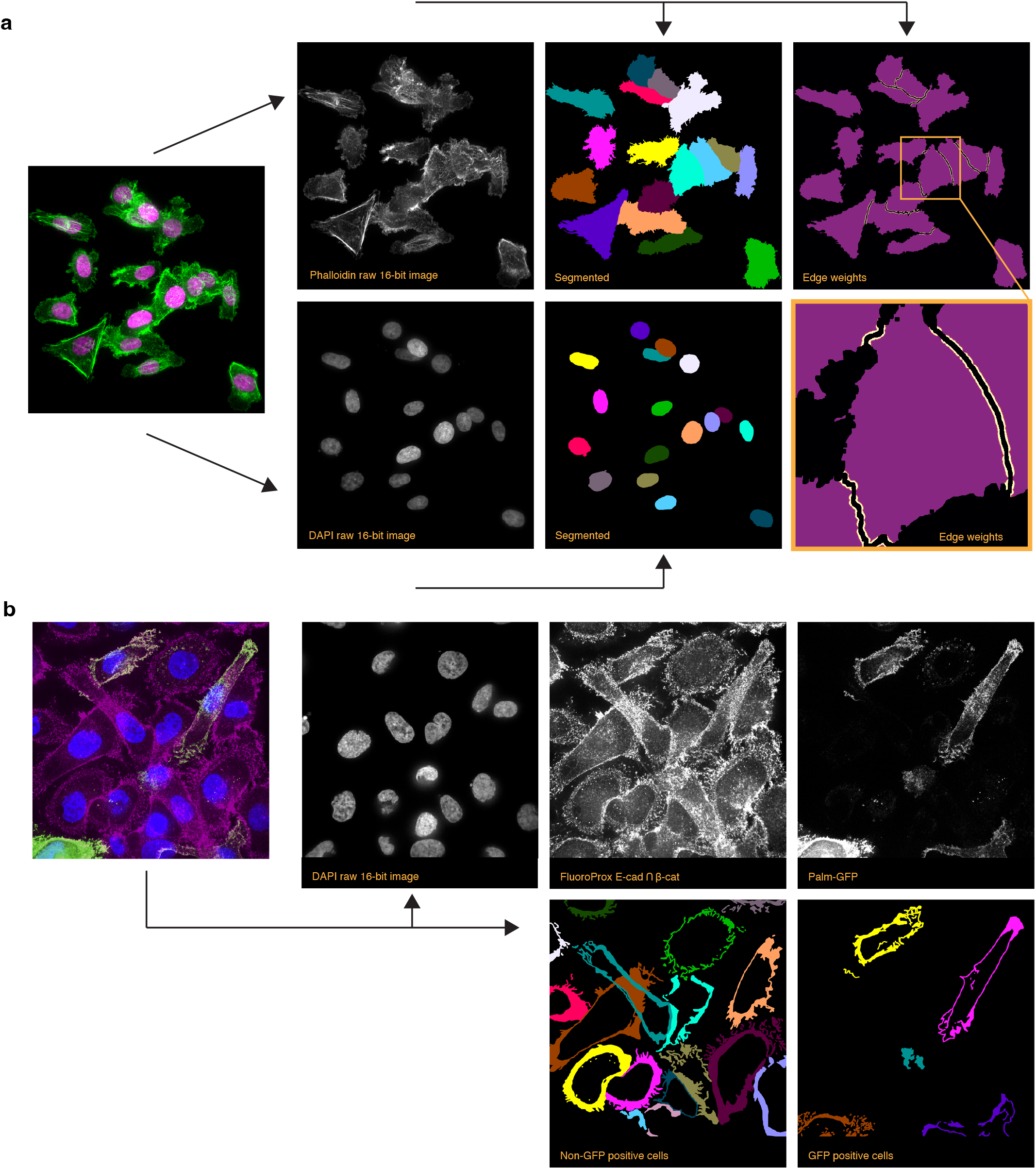
Segmentation of cellular components. **a)** Segmentation of cytosol and nucleus from phalloidin and DAPI stains, respectively. The leftmost panel shows a composite image of cells stained with phalloidin (green) to label actin filaments and DAPI (magenta) to label nuclei. The far-right panel shows the edge weights used to refine the segmentation of cytosol signals. **b)** Segmentation of cell membranes from Palm-GFP positive and negative cells. The leftmost panel displays a composite image, where the cell membranes are stained with FluoroProx (to label E-cadherin and β-catenin) in magenta and Palm-GFP in green and cell nuclei (DAPI) in blue. Segmented membrane masks are then used to measure the average fluorescent signal at the membrane.

**Figure S4.**
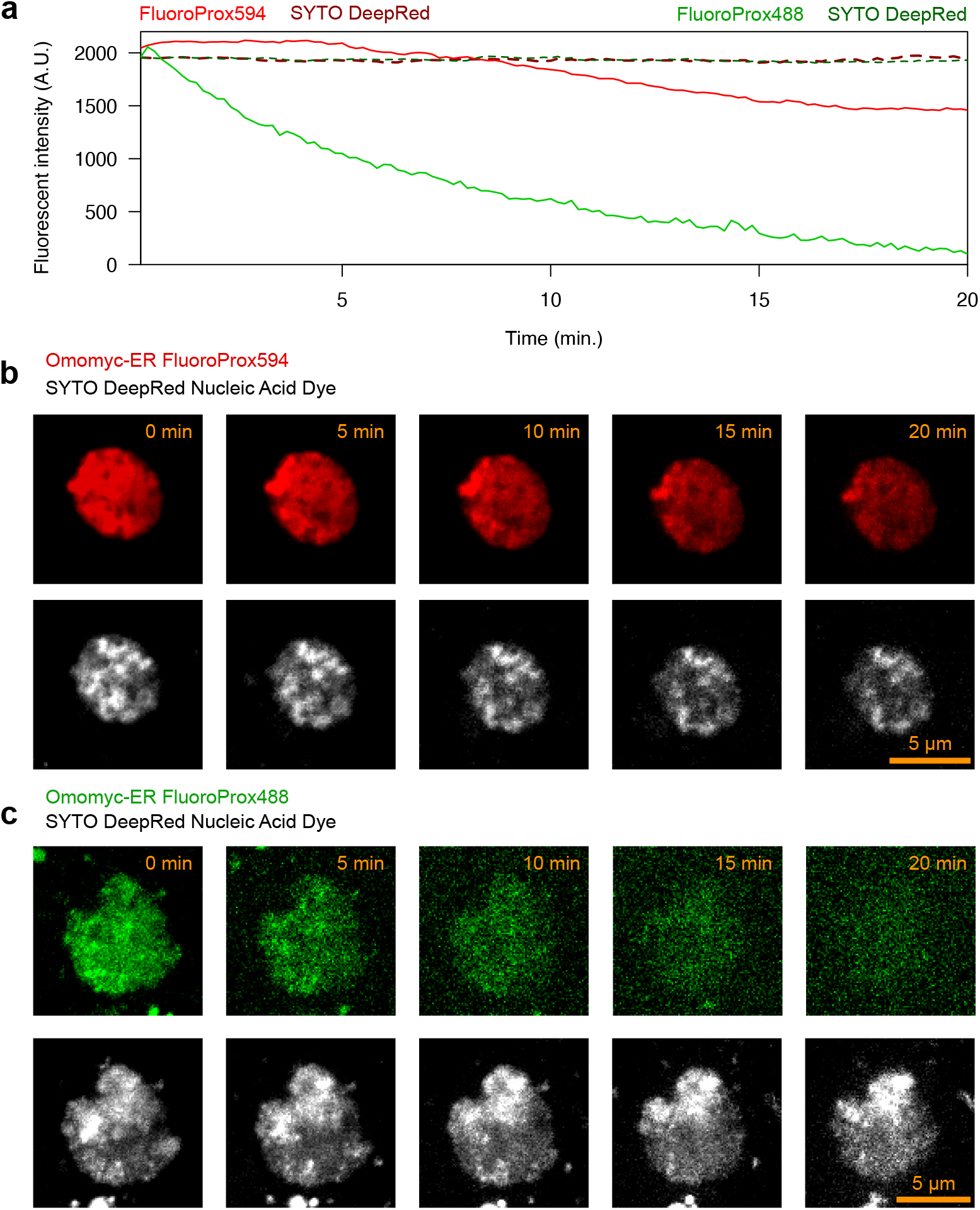
Comparative photostability among FluoroProx probes in living cells. While both FluoroProx594 and FluoroProx488 work fine in fixed cell assays using antibodies, in living cell condition FluoroProx488 is more succesible to oxidation and subsequent bleaching. A fact that has to be taken into consideration when designing experiments. **a)** Average fluorescent signal over time. Images taken every ten seconds (0.1 Hz). Darker dashed lines indicate counter staining for DNA using SYTO DeepRed Nucleic Acid Dye. Red lines shows signals from FluoroProx594 experiment and green lines show signals from FluoroProx488. **b)** Example of FluoroProx594 (red) and SYTO DeepRed nucleic acid dye (DNA) in a single U2OS cell. **c)** U2OS cell example of FluoroProx488 (green) and DNA stained as in b). Scale bars: 5 µm.

